# Sensitivity to perturbations of a cell migration under temporal regulation

**DOI:** 10.1101/2020.07.21.213710

**Authors:** Clément Dubois, Shivam Gupta, Andrew Mugler, Marie-Anne Félix

## Abstract

Few studies have measured the robustness to perturbations of the final position of a long-range migrating cell. In the nematode *Caenorhabditis elegans*, the QR neuroblast migrates anteriorly in the young larva, while undergoing three rounds of division. The daughters of QR.pa stop their migration at an anterior body position and acquire a neuronal fate. Previous studies showed that the migration stops upon expression of the Wnt receptor MIG-1, which surprisingly is not induced by positional cues but by a timing mechanism (Mentink et al. 2014). Given this temporal regulation, we wondered 1) how precise QR.pax positioning is when confronted with various challenges, such as stochastic noise, environment or body size variation and 2) whether QR.pax position varies among *C. elegans* wild isolates. We find that the variance of QR.pax final position is similar to that of other long-range migrating neurons. Its mean position undergoes a slight posterior shift at higher temperature, while its variance is greatly increased following sustained starvation at hatching. We manipulated body size using mutants and tetraploid animals. As expected from the temporal mechanism, smaller mutants display anteriorly shifted QR.pax cells, while longer mutants and tetraploids display posteriorly shifted QR.pax cells. Using a mathematical model, we show however that body size variation is partially compensated. We find that cell speed is indeed altered in body size mutants. Finally, we could detect highly significant variation among *C. elegans* wild isolates. Overall, this study reveals that the final cell position of QR.pax shows some degree of sensitivity to external perturbations and natural genetic variation.

## Introduction

Cell migration is a key process during the development of many animal tissues. Much is known about the cues that confer directionality to the migration (Whangbo and Kenyon, 1999; Branda and Stern, 2000; Duchek et al., 2001; Pani and Goldstein, 2018; Szabó and Mayor, 2018; Kim et al., 2019), accuracy in gradient sensing and directionality (Barkai and Leibler, 1997; Endres and Wingreen, 2008) and the signal transduction and cytoskeletal dynamics behind the movement (Abercrombie et al., 1970; Ridley et al., 2003; Svitkina, 2018; van Helvert et al., 2018). In comparison, the termination of migration has been little studied, although it is obviously of key importance for final cell and organ position (Meighan and Schwarzbauer, 2007; Aman and Piotrowski, 2010; Inamura et al., 2012; Kikuchi et al., 2015; Paksa et al., 2016). Termination of most cell migrations is thought to be spatially triggered by homogenous concentrations of the guiding cues (Doitsidou et al., 2002), adhesion to a specific target cell (Rohrschneider and Nance, 2013) and/or physical barriers (Halfter et al., 2002; Paksa et al., 2016), but in a recent case was shown to be regulated independently of spatial cues, in a time-dependent fashion (Mentink et al., 2014, see below).

The occurrence of cell migration begs the question of the degree of precision in final cell position. The degree of sensitivity of a trait to a given perturbation — or conversely its precision or robustness — is a fundamental characteristic of biological systems (Félix and Wagner, 2008). Although defects in the direction of cell migration are often reported (Burke et al., 2015), few studies have measured the precision in the length of migration and the final position of the cell (Branda and Stern, 2000; Grimbert et al., 2016; Paksa et al., 2016; Lau et al., 2020). Moreover, while the majority of cell migration studies explore the effect of genetic or experimental perturbations, they have not addressed more ecologically and evolutionary relevant types of perturbation, such as environmental, stochastic, or natural genetic variation.

The QR neuroblast is a cell that migrates a long distance from the posterior to a more anterior position during the first larval stage of the nematode *Caenorhabditis elegans.* Three rounds of QR cell division take place during or at the end of the migration (Sulston and Horvitz, 1977). The progeny are named according to their anterior or posterior position at each successive cytokinesis: thus QR.p is the posterior daughter of QR, and QR.pa the anterior daughter of QR.p. Finally, the daughter cells of QR.pa, called QR.paa and QR.pap (hereafter called QR.pax), acquire a neuronal identity (Chalfie and Sulston, 1981; White et al., 1986). Much is known about the signaling pathways, transcription factors and cytoskeleton regulating their posterior-to-anterior direction and orientation of migration (Middelkoop and Korswagen, 2014; Josephson et al., 2016; Rella et al., 2016). Concerning the termination of migration and final cell positioning, the posterior-to-anterior QR.pa lineage stops upon expression of the Wnt receptor MIG-1 (Mentink et al., 2014). Surprisingly, the expression of *mig-1* in QR.pa is not induced by the cell reaching a certain position in the body, but by a temporal regulation, independently of cell position. Indeed, preventing QR migration or increasing its speed does not alter the timing of *mig-1* expression (Mentink et al., 2014). After QR.pa stops migrating, its daughter cells QR.pax separate in a dorso-ventral direction while crossing each other in an antero-posterior direction (Rella et al., 2016; Altun and Hall, 2020); they then differentiate without further change in cell body position (Figure 1).

**Figure 1:**
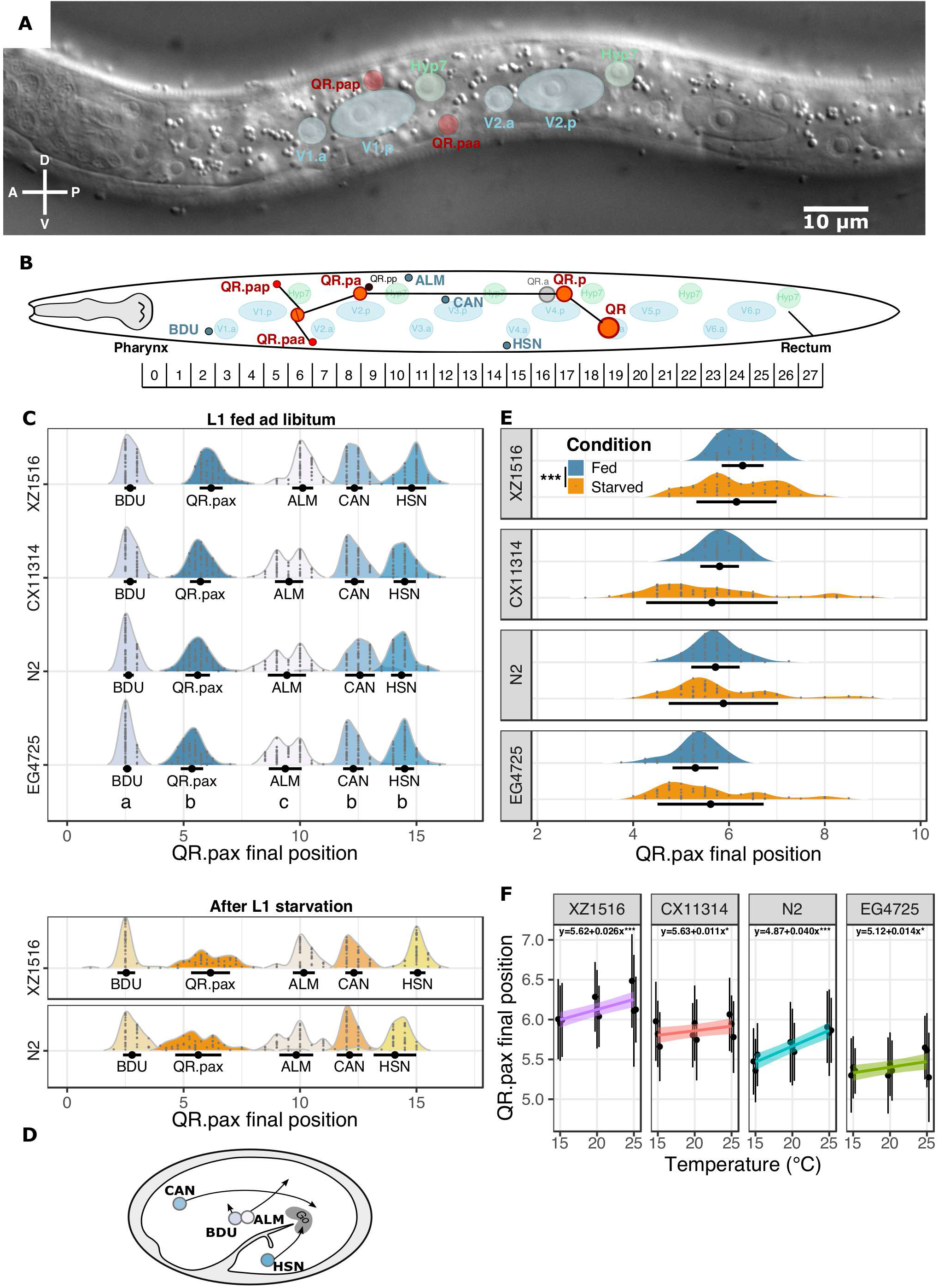
QR.p neuroblast migration during the L1 stage and its sensitivity to noise and environmental variations. **A)** Nomarski micrograph of a late L1 larva, where QR.pap and QR.paa have reached their final position. The relevant cells are outlined. **B)** Illustration of QR.p neuroblast migration during the L1 stage showing the final position of QR.paa and QR.pap and the relative scale used in scoring their position. This relative scale from 0 to 27 was constructed relative to the seam cells, as shown here. **C)** Sensitivity of QR.pax final position to noise. We measured the final position of QR.pax and that of other neurons that migrate, albeit on shorter distances during embryogenesis: BDU, ALM, CAN and HSN. Black dots and error bars represent the mean and standard deviation (SD), respectively; n=70 per strain, 2 replicates merged. Letters a,b,c represent groups of cells with similar variance (Levene’s test for homogeneity of variance). The two bottom panels show final position between neuron positions after early L1 starvation. **D)** Illustration of BDU, ALM, CAN and HSN migration during embryogenesis, adapted from Altun and Hall (2020). **E)** Sensitivity of QR.pax to a starvation treatment in the L1 stage. After food depletion, arrested larvae were kept starved for two days before being fed again and allowed to develop. Black dots and error bars represent the mean and SD; n≥40 per strain and condition. (Levene’s test for homogeneity of variance, *** p-value < 0.001). **F)** Robustness of QR.pax to temperature. Plates containing L1 larvae were transferred from 20°C to different temperatures one generation before scoring. Correlation between QR.pax final position and the growth temperature of different wild genotypes, in three independent replicate experiments. Black dots and error bars represent the mean and SD of each replicate, the colored lines represent the regression line with the associated equation estimated with a linear mixed model. (n≥40 per strain, temperature and replicate, ***: *p*<0.001, *: *p*<0.05).

We here probe the sensitivity of this system to various perturbations of ecological and evolutionary relevance. We score both the mean and the distribution of the final cell position.

First, noise, or stochastic variation, is measured by variation in the trait among isogenic animals grown in the same environment (Spudich and Koshland, 1976; McAdams and Arkin, 1997; Elowitz et al., 2002). In our study, we measured the degree of robustness (or sensitivity) of QR.pax final position by comparing its variance to that of other migrating neuronal cell bodies: those of ALMR, BDUR, CANR and HSNR (later called ALM, BDU, CAN and HSN) (Sulston et al., 1983; Hedgecock et al., 1987) (Figure 1.D).

Second, we perturbed the external environment in two ways. We first tested the effect of an environmental perturbation that is highly relevant in the wild, namely different temperatures, from 15°C to 25°C. Second, we tested the effect of starvation at hatching, before QR migration. After embryonic development inside its egg shell, *C. elegans* hatches in the first larval stage (L1) in the shape of a small worm. If no food is provided, the larva enters a developmental arrest program that can last for several days, prior to QR migration and first division (Johnson et al., 1984; Baugh, 2013). QR migration starts shortly after feeding the animal. We thus tested the robustness of QR.pa final position to this temporary developmental arrest after hatching.

Third, we reasoned that given the temporal regulation of *mig-1* expression and of the migration endpoint, a change in L1 larva body size would lead to a shift in the final cell position relative to other landmarks along the antero-posterior axis — provided that cell migration occurs at the same speed in animals of different body sizes. The expectation is that in longer animals QR.pa stops in a more posterior position relatively to body landmarks; and in smaller animals at a more anterior position. To address the effect of body size on the relative position of QR.pax, we used different body size mutants in the reference genetic background N2. In parallel, we developed a mathematical model of the expected relationship between body size and QR.pax final position, taking into account larval growth during cell migration.

Fourth, we explored natural variation using a panel of *C. elegans* wild isolates. The study of wild isolates reveals natural variation for phenotypic traits (Hodgkin and Doniach, 1997; Farhadifar et al., 2015; Cook et al., 2016; Gimond et al., 2019; Lee et al., 2019) and different genotypes can reveal different sensitivities to perturbations (Braendle and Félix, 2008). So far, QR.pax final position had only been measured in the laboratory-evolved reference background N2 and mutants from this strain. Laboratory adaptation may alter many phenotypes in many organisms including *Drosophila melanogaster* (Stanley and Kulathinal, 2016), *Caulobacter crescentus* (Marks et al., 2010) or *Caenorhabditis elegans* (McGrath et al., 2009; Sterken et al., 2015). We therefore wondered to which extent the final position of QR.pax evolved within *C. elegans*; and whether the degree of robustness to noise and environmental perturbations was a shared feature between laboratory adapted genotypes and wild isolates.

We find that the variance of QR.pax final position is similar to that of other neurons that migrate at a long-range during embryogenesis. At higher temperatures, the mean position of QR.pax undergoes a slight posteriorly shift, while its variance is increased by a starvation treatment after hatching, showing that the system is not fully robust to external perturbations. Moreover, as expected from the temporal mechanism of migration termination, smaller mutants display more anterior QR.pax cells, while longer mutants and tetraploids display posteriorly shifted QR.pax cells. Finally, we could detect highly significant variation among *C. elegans* wild isolates that was not significantly explained in our tested panel by variation in body size.

## Materials & methods

### *C. elegans* strains

*Caenorhabditis elegans* were grown at 20°C on 55 mm diameter Petri dishes with NGM, fed on *Escherichia coli* OP50 according to the standard procedures (Brenner, 1974). To investigate the relationship between body size and QR.pax final position, we selected a list of mutations in the reference genetic background N2 known to affect egg size and a priori not cell migration: JJ1271 *glo-1(zu391)* (Hermann et al., 2005), GL302 *cid-1(rf34)* (Olsen et al., 2006), RB2488 *amph-1(ok3443)* (The C. elegans Deletion Mutant Consortium, 2012), RB1658 *mca-3(ok2048)* (The C. elegans Deletion Mutant Consortium, 2012) BA819 *spe-11(hc77)* (L’Hernault et al., 1988), GH383 *glo-3(zu446)* (Rabbitts et al., 2008), JT513 *nrf-5(sa513)* (Choy and Thomas, 1999), JT73 *itr-1(sa73)* (Iwasaki et al., 1995), RT362 *rme-4(b1001)* (Sato et al., 2008), JK4545 *larp-1(q783)* (Nykamp et al., 2008), CB30 *sma-1(e30)* (Brenner, 1974), RB1977 *fln-1(ok2611)* (The C. elegans Deletion Mutant Consortium, 2012), CB185 *lon-1(e185)* (Brenner, 1974) and the tetraploid 4A:4X strain SP346 (Madl and Herman, 1979).

To investigate the natural variation for the trait, we used the following wild isolate strains (Cook et al., 2017) CB4856, CB4932, CX11271, CX11314, ECA189, ECA191, ECA36, ECA363, ECA369, ECA372, ECA396, EG4725, JT11398, JU1200, JU1242, JU258, JU363, JU367, JU775, JU830, JU1393, LKC34, MY10, NIC199, NIC251, NIC256, NIC268, PB303, PB306, QG2075, QG556, QX1791, QX1793, QX1794, WN2001, XZ1513, XZ1514, XZ1515, XZ1516.

The strain NH646 *ayIs9[Pegl-17::gfp + dpy-20(+)]; dpy-20(e1282ts) (Branda and Stern, 2000)* with a Q lineage GFP reporter was also used in this study (kind gift of the Korswagen laboratory). The strain GOU174 *casIs35*[*Pgcy-32*::mCherry, *unc-76(+)*] X; *zdIs5*[*Pmec-4*::gfp, *lin-15(+)*]I (Zhu et al., 2016) was used to observe cell body and axon of mechanosensory neurons including AVM (QR.paa) (kind gift from the Ou laboratory).

### QR.pax final position measurements in N2 mutants and wild isolates

QR.paa and QR.pap final positions were measured using Nomarski microscopy of late L1 larval stage animals mounted on 3% agar pads and immobilized with 1mM sodium azide. At this stage, the V seam cells have divided once, QR.paa has reached its final ventral position and QR.pap its final dorsal position in the animal. QR.pax final position is the mean position of QR.pap and QR.paa in each individual, measured with a relative scale based on the V-derived seam cells (Harris et al., 1996; Whangbo and Kenyon, 1999; Coudreuse et al., 2006; Mentink et al., 2014). Note that the relative position of QR.paa to QR.pap does not change when the long-range migration of their ancestors is affected (Fig. S3 in Mentink et al., 2014). We constructed a semi-discrete scale from the pharynx (0) to the rectum (27), based on the repeated pattern formed by the nuclei of the seam cells and hyp7 cells. A half value is attributed when the nucleus is between two landmarks. In some cases, we also measured distances in microns.

### Robustness to noise

We measured the variance of QR.pax position when faced with stochastic noise and compared it to that of other neurons migrating during embryogenesis: BDU, ALM, CAN and HSN (Sulston et al., 1983; Hedgecock et al., 1987). These neurons are easily recognizable and are in the same environment as QR.pax. We could use the same relative scale to measure the position of these cells’ bodies and compare variance between them. We performed a Levene’s test for equality of variance to measure the differences of variance between strains and cell positions, adjusted with Bonferroni correction for multiple testing. The experiment was performed in two replicates and those were merged as there was no replicate effect on the variance (see Supp. Table 1).

### Robustness to environmental perturbations

In order to test the robustness of QR.pax final position to growth at different temperatures, L1 larvae from the N2 strain and three wild isolates were transferred from 20°C to 15°C, 20°C or 25°C one generation before scoring. We repeated this experiment three independent times. We performed a two-way ANOVA to explain QR.pax final position according to temperature of growth (considered as a continuous variable), strain, temperature x strain interaction and replicate experiments. The effect of the replicates adds variability in the measurements but not in a particular direction (see Supp. Table 1). To consider the variability in the response to temperature, we modeled the replicate effect as a random effect. We tested the correlation between QR.pax final position and the absolute value of the latitude of the wild isolate sampling location (as a proxy for the temperature of origin), using Pearson’s product-moment correlation test.

To test the robustness to transient developmental arrest in the L1 stage, cultures were grown until food depletion. Once food was exhausted on the plate, we maintained the population under starvation for two more days. Note that this protocol avoided a bleach treatment that may affect the larvae independently of the starvation period. Then, we collected larvae in M9 buffer and plated them onto NGM agar plates with *E. coli* OP50. We scored QR.pax position six to eight hours later, only when V seam cells divided once, Pn cells migrated ventrally, QR.pap reached the dorsal part of the animal and QR.paa the ventral part. We thus did not score the most delayed animals – note that rejecting these animals is unlikely the reason for the observed increased variance in QR.pax position. In contrast to the bleaching method, small larvae may have access to a minimal amount of food before starvation allowing the beginning of QR migration. We found that using this protocol, 58% of animals have QR.pa divided above the seam cell V1, 24% with QR.pa at the top of V2, 12% at the top of V3 and 6% at V4 (n=50 worms). We performed a Levene’s test for equality of variances to measure the effect of the strain and the starvation condition on QR.pax final position variance, adjusted with Bonferroni correction for multiple testing. During this experiment, the position of BDU, ALM, CAN, HSN and QR.pax was measured in XZ1516 in parallel. The same method was then used in an independent experiment using the genotype N2.

### Distance measurements

We first measured egg length in a set of mutants with a putatively altered embryo size (see list above) and compared their size to that of N2 using a two-sided t-test with Bonferroni adjustment of the *p*-value for multiple comparisons. Most body size mutants concern larval growth and not earlier stages. Live embryos of *cid-1(rf34), glo-3(zu446), itr-1(sa73), mca-3(ok2048), rme-4(b1001), spe-11(hc77)* mutants did not show significant differences in length compared to N2 and were thus rejected (Figure S2A).

We measured QR.pax final position in the remaining mutants with significant differences in egg size. Three mutants had gross defects and we decided to discard them further from the analysis: *amph-1(ok3443)* and *nrf-5(sa513)* exhibited several defects including cell migration defects for BDU, ALM and/or dorsal-ventral migration of QR.paa and QR.pap; *glo-1(zu391)* mutants had long eggs but small and sick larvae. We then compared QR.pax final position of the mutants to N2 with a two-sided t.test and p-values were adjusted with Bonferroni method for multiple comparisons.

To test the correlation between egg size and L1 larva body size, we measured the distance between the rectum and the end of the pharynx, corresponding to the area of migration of the QR neuroblast from 27 to 0 in our relative scale. Plates containing eggs were washed several times and transferred onto a fresh plate in order to only keep unhatched embryos. Freshly hatched larvae (0-20 min) were mounted on agar pads immobilized with 1mM sodium azide and measured from the pharynx to the rectum, or transferred to fresh plates to perform the measurements 6h after hatching.

To test whether body size differentially affected each step of the migration, we measured the distance between the rectum, QR.pp (QR.p site of division), QR.pa and the pharynx, in worms where QR.pa had just divided and the QR.pp apoptotic body was still present. To visualize the cells in body size mutants, we introgressed *ayIs9[Pegl-17::gfp,dpy-20+]* in *sma-1(e30)* and *lon-1(e185)* background by crossing CB30 and CB185 strains, respectively, with NH646. The strains of genotype *ayIs9 sma-1(e30)* and *lon-1(e185); ayIs9* were used to estimate cell velocity of QR by measuring the distance between the rectum, the QR lineage and the pharynx at 3h and 6h after hatching, corresponding to QR.p lineage long-range migration.

To investigate the effect of QR.pax final position on neuronal differentiation phenotypes, we applied the starvation protocol described above to the strain GOU174 wiht a GFP marker for mechanosensory neurons and monitored the axons of AVM (QR.paa), PVM, ALM and PLM. We measured the relative position of AVM by dividing the Pharynx-to-AVM distance by the Pharynx-to-ALM distance. We measured AVM position in the L3 stage, 31 hours after food provisioning to starved worms, with two independent replicate experiments. The control population was obtained by washing plates with M9 to only keep embryos on NGM. Then the embryos were allowed to hatch and feed on OP50 for 41 hours before phenotyping. We performed a Levene’s test for equality of variances to measure the effect of starvation on the variance in AVM relative position in each replicate and in the whole dataset. The effect of starvation and AVM final position on the axon overlapping phenotype was analyzed with a generalized mixed model.

Pictures and measurements were performed with the camera Photometrics CoolSNAP ES (Roper Scientific) and software Nikon NIS element D (version 3.1). The size in pixel was then converted to micrometers after calibration (Size(μm) = Size(px)/9.84).

### Relationship between QR.pax final position and distance measurements

We used the data of egg length, pharynx-to-rectum distance (hereafter called P-to-R) at two timepoints (0 and 6 hrs) and QR.pax final position of each strain to test relationships between these variables. The different size measurements and QR.pax final positioning was not measured in the same individual but in an isogenic population. To consider the variance in both axes for the regressions we used a bootstrapping approach with 1000 iterations. Each iteration was made by subsampling 20 egg size measurements, P-to-R distance at hatching, 6 hours after hatching and QR.pax final position per genotype. Model 1 regression (Ordinary Least Square method) was performed between each variable at each iteration. Means, intercepts, slopes, *R*^*2*^_*adj*_ and *p*-values generated per iteration were saved for plotting. The median of the bootstrapped *R*^*2*^_*adj*_ and *p*-values of the regression was used to conclude on the likelihood of the correlation.

### Mathematical model for QR.pax final position

For the no-compensation model, we treated the dynamics of the QR cell lineage as one-dimensional, constant-velocity migration within a growing larva. Taking the pharynx to be stationary at the origin *x* = 0 and the rectum to be moving away with constant velocity *u* due to the growth, the length of the pharynx-to-rectum region evolves in time according to

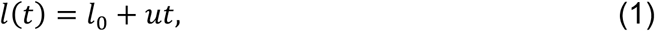

where *l*_0_ is the length at hatching (*t* = 0). The dynamics of the cell position *x* are

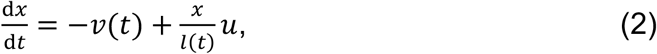

where the first term is the leftward velocity of the cell, and the second term is an effective rightward velocity due to the growth. In the second term we enforce uniform expansion of the larva during growth, such that intermediate points move according to the fraction of the distance to the origin. We assume that the cell begins migrating at a constant velocity *v*_0_ at a time *t* = *τ* after hatching and stops migrating at a time *t* = *T*,

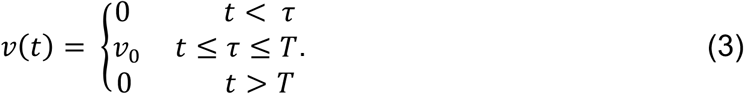

Integrating Eq. 2 gives

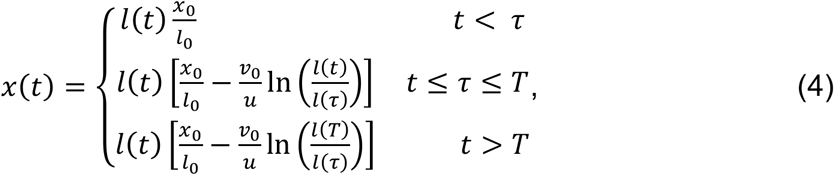

where *x*_0_ is the initial position of the cell. The relative position on the semi-discrete scale is *p* = *Nx*/*l*, where *N* = 27 (Figure 1B). Inserting Eq. 4, we have

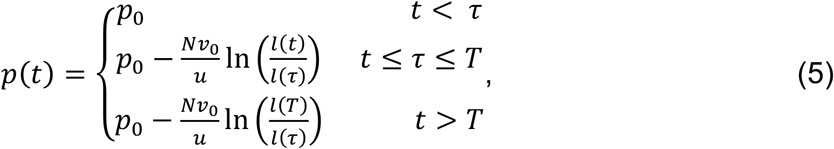

where *p*_0_ = 19 is the initial position of the cell (Figure 1B). We take *u* = (35 μm)/(6 hours) = 5.8 μm/hr from the experiments (Figure S2E), and *τ* = 2 hr and *T* = 8 hr (Sulston and Horvitz, 1977; Ou and Vale, 2009; Ebbing et al., 2019) from the literature, leaving only *v*_0_ as a fit parameter. Figure 2C (red curve) shows *p* versus *l*_0_ from Eq. 5 (*t* > *T*) with best-fit value *v*_0_ = 11.6 μm/hr.

**Figure 2:**
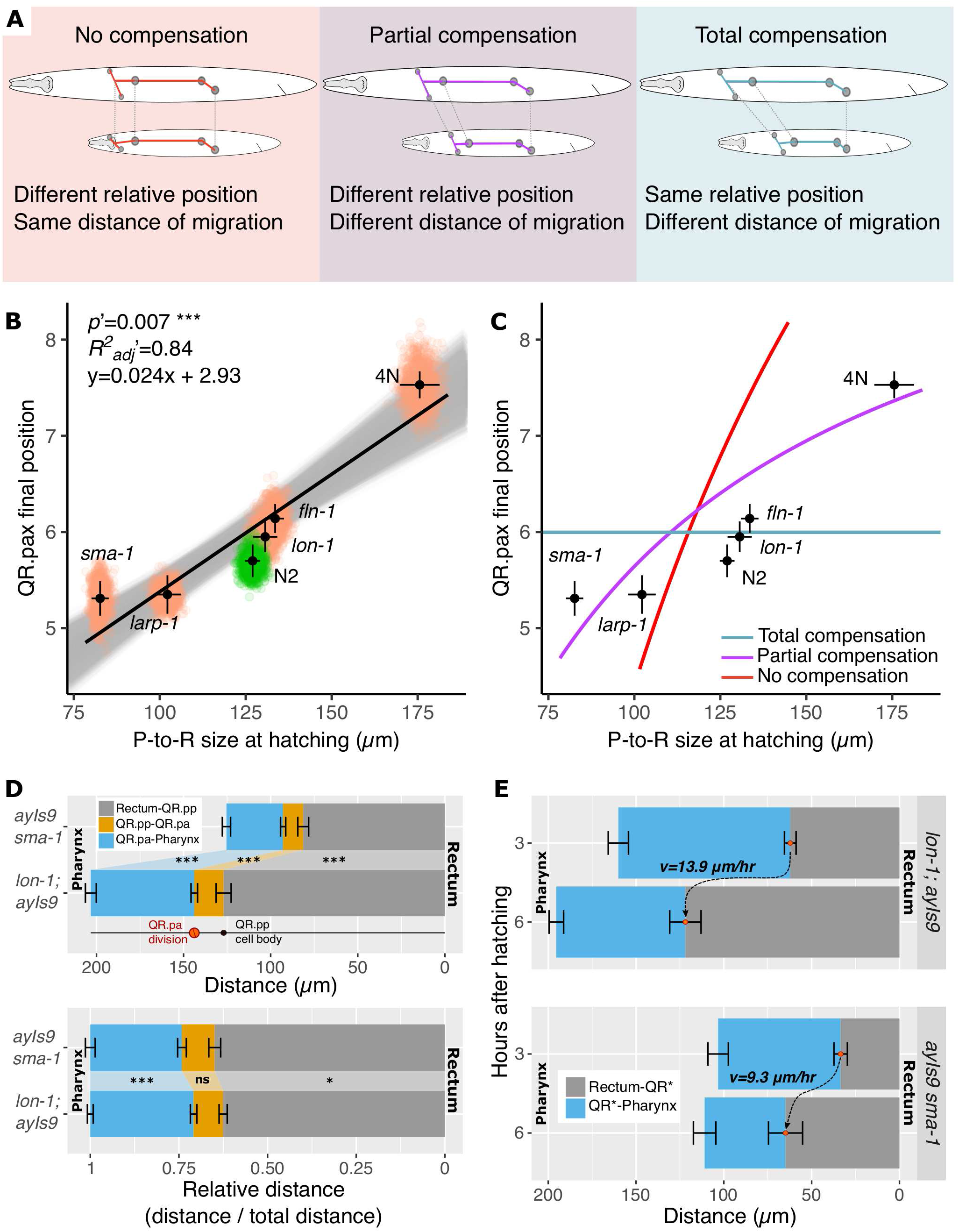
Sensitivity of QR.pax position to body size. **A)** Schematics of the relative position of QR.pax in a long versus a short animal in the absence of body size compensation (left), with full compensation (right) or with a partial compensation (middle). **B)** Relationship between QR.pax final position and Pharynx-to-Rectum distance at hatching relationship in a subset of the panel. Orange and green dots represent one round of subsampling of the data. Black dots and error bars represent the mean and confidence intervals (CI 95%) from the data for each genotype. The black line represents the regression line from the data. The grey area represents regression lines after each iteration of subsampling. **C)** Mathematical model of the relationship between QR.pax final position and Pharynx-to-Rectum distance assuming full body size compensation, no compensation (with one fit parameter, cell velocity), or partial compensation (with no fit parameters). **D)** Location of QR.p and QR.pa divisions in *ayIs9 sma-1* and *lon-1; ayIs9* backgrounds, in absolute value of distance to the rectum and pharynx or scaled relatively to body size **(E)**. The apoptotic body of the QR.pp cell marks the site of QR.p division. Bars and error bars represent the mean and confidence intervals (95%) per genotype; n>20 per strain. Two-sided t-test: ***: *p*<0.001, *: *p* <0.05, *n.s.*: not significant. **F**) Absolute distances between the rectum, QR cell* and the pharynx 3h and 6h after hatching in *lon-1; ayIs9* and *ayIs9 sma-1* animals. At 3h QR* is either QR or QR.p in some animals, at 6h QR* is either QR.p division or QR.pa in *lon-1; ayIs9* and mostly QR.p in *ayIs9 sma-1*. Bars and error bars represent the mean and confidence intervals (95%) per genotype; n≥10 per genotype. The cell velocity *v* was calculated with the Eq. 9.

For the partial-compensation model, we used Eq. 4 to infer the cell velocity from measurements of *x* and *l* at two time points *t*_1_ = 3 hr and *t*_2_ = 6 hr. Calling *x*(*t*_1_) = *x*_1_, *x*(*t*_2_) = *x*_2_, *l*(*t*_1_) = *l*_1_, and *l*(*t*_2_) = *l*_2_ for short, and recognizing that both *t*_1_ and *t*_2_ fall between *τ* and *T*, we have from Eq. 4,

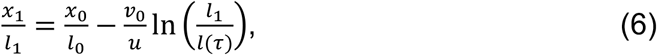

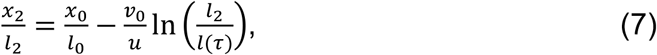

at each time point. Subtracting Eq. 7 from Eq. 6 gives

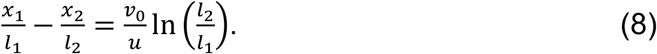

Because growth is linear in time (Eq. 1), we have *u* = (*l*_2_-*l*_1_)/(*t*_2_-*t*_1_). Inserting this expression into Eq. 8 and solving for *v*_0_ gives

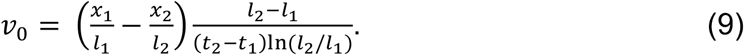

Eq. 9 was used to calculate the cell velocities in the *lon-1(e185)* (*x*_1_ = 97.9 μm, *x*_2_ = 73.5 μm, *l*_1_ = 160.1 μm, *l*_2_ = 195.4 μm) and *sma-1(e30)* (*x*_1_ = 69.8 μm, *x*_2_ = 46.1 μm, *l*_1_ = 103.2 μm, *l*_2_ = 110.9 μm) mutants (Figure 2E), giving *v*_0_ = 13.9 μm/hr and *v*_0_ = 9.3 μm/hr, respectively. Assuming a linear relationship between velocity and body size,

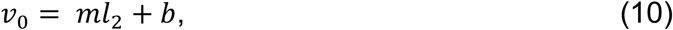

the two values of *v*_0_ and *l*_2_ imply *m* = 0.054 hr^−1^ and *b* = 3.3 μm/hr. Recognizing that *l*_2_ = *l*_0_ + *ut*_2_, Eq. 10 becomes

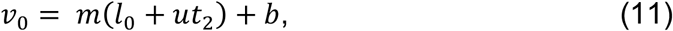

with *ut*_2_ = 35 μm (Figure S2E). Figure 2C (purple curve) shows *p* versus *l*_0_ from Eq. 5 (*t* > *T*) with Eq. 11 inserted for *v*_0_.

### Statistical analyses, plots and raw data

Statistical analyses and plots were performed using R version 3.5.2 (R core Team, 2018), R studio version 1.1.463 (RStudio Team, 2015) and the following packages: car (Fox and Weisberg, 2019), lmerTest (Kuznetsova et al., 2017), Rmisc (Hope, 2013), ggplot2 (Wickham, 2016), ggstance (Henry et al., 2019), ggrepel (Slowikowski, 2019). The distribution of QR.pax final position in Figure 1, 3A and Figure S2B was represented with ggridges (Wilke, 2018) using a bandwidth of 0.25 for smoothing. Raw data and summary are presented in the Supplementary Table.

**Figure 3:**
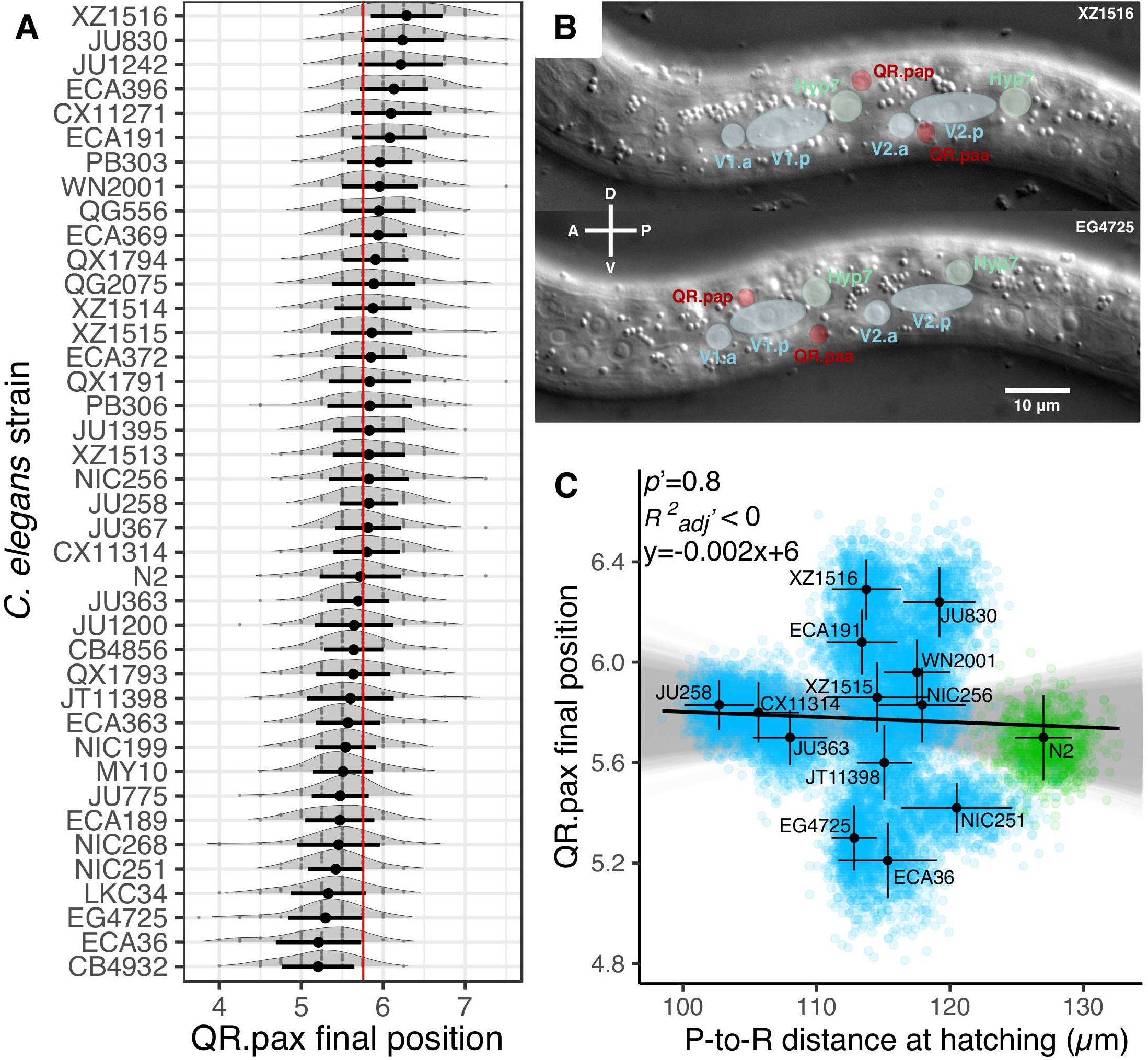
Natural variation of QR.pax final position in *C. elegans*. **A)** Natural variation of QR.pax final position in a panel of 40 *C. elegans* wild isolates. Grey dots represent QR.pax final position for each animal; black dots and error bars represent the mean and SD per genotype, respectively (n=50 per strain). The red dashed line indicates the grand mean over the 40 isolates. **B)** Nomarski micrograph of late L1 larvae showing an example of posterior (strain XZ1516, top) and an anterior (strain EG4725, bottom) QR.pax relative position. **C)** Relationship between QR.pax final position and Pharynx-to-Rectum distance at hatching in a subset of the panel. Blue and green dots represent the mean after one round of subsampling. Black dots and error bars represent the mean and CI (95%) for each genotype from the data. The black line represents the regression line from the data. The grey area represents regression lines after each iteration of subsampling.

## Results

### Variance in QR.pax final position is similar to that of other migrating neurons

The sensitivity to noise, or stochastic variation, is measured by the variance of a trait in a population of isogenic individuals in a constant environment. We first compared in N2 and three genetically distinct wild isolates the variance in QR.pax final position to that of other neurons that migrate during embryogenesis: BDU, ALM, CAN and HSN (Sulston et al., 1983; Hedgecock et al., 1987) (Figure 1C,D). The cell body position of these neurons can be measured in the same animals with the same scale as that of QR.pax (Harris et al., 1996; Forrester and Garriga, 1997; Ch’ng et al., 2003; Zinovyeva and Forrester, 2005; Zinovyeva et al., 2008). QR.pax final position variance is similar within the four strains (*F*=0.48, *p*=1) and the two replicates (*F*=0.15, *p*=1) but not among the six neurons (*F*=37.8, *p*<10^−15^). BDU has significantly lower variance than the other migrating neurons, which is likely to reflect the fact that it only migrates at a short range (Figure 1D). The variance of QR.pax is similar to those of CAN (*F*=5.5, *p*=0.19) and HSN (*F*=4.4, *p*=0.36). ALM has a slightly larger variance than the other neurons (*F*>9, *p*<0.02). Altogether, this result reveals that QR.pax final position is quite robust to noise despite the temporal regulation of its migration endpoint. Note that among wild isolates, we do not observe consistent co-variation between migrating neurons except for QR.paa and QR.pap (Suppl. Table). This also imply that using QR.pax as the mean position between QR.paa and QR.pap is relevant.

### Temperature mildly affects the mean position of QR.pax, while early developmental arrest affects its variance

We tested two ecologically relevant environmental variables for their effect on QR.pax positioning: temperature and starvation at hatching. Temperature can affect various developmental processes. Plates containing L1 larvae were transferred from 20°C to different temperatures (15°C, 20°C or 25°C) one generation before scoring. The same four strains were scored as above. The major effect on the final positioning is due to the genotype (*X*^2^=568.67, *p*<10^−15^). The temperature shifts QR.pax final position (*X*^2^=66.93, *p*<10^−15^) but the strength of this effect is slightly modulated by the genotype (genotype x temperature interaction: *X*^2^=16.23, *p*=0.001). The temperature shifts QR.pax final position toward the anterior as the temperature decreased, with a value of 0.04 relative position unit per degree (Figure 1F). This value decreases to 0.03 in the genotype XZ1516, 0.01 in the genotype EG4725 and CX11314, suggesting a lesser sensitivity to temperature in the two last wild isolates. Note that temperature is not a specific environmental variable as it may affect many processes in the larva.

After hatching in the absence of food, larvae of *Caenorhabditis elegans* stop their development, a phenomenon known as the L1 arrest, regulated by insulin signaling (Kaplan and Baugh, 2016; Zheng et al., 2018). Development starts again when the larvae ingest food, but various developmental processes may not be tightly synchronized and many traits may be affected (Lee et al., 2012). We maintained recently hatched larvae under food deprivation for two days in crowded plates. We then provided them with food and scored them six to eight hours later, once QR.paa had reached the ventral part of the animal, QR.pap the dorsal part (and thus both had stopped their migration) and the V seam cells had divided once. The variance among the different fed strains was similar (*F*=0.48, *p*=1) as well as the variance among the starved strains (*F*=2.1, *p*=0.30). Nonetheless, starvation increased considerably the variance in the starved treatment compared to continuously fed worms (*F*=13.37, *p*<10^−14^) (Figure 1E). Note that after L1 starvation, the variance of other neurons does not seem to be largely altered, especially in XZ1516 (Figure 1C).

Differences in final cell position may result in downstream differentiation phenotypes, such as variation in axon morphology in the case of neurons. QR.paa becomes the AVM mechanosensory neuron. We thus monitored axons of mechanosensory neurons after starvation of GOU174, a strain exhibiting a GFP reporter in mechanosensory neurons including AVM (see Methods). We scored larvae at the L3 stage and used the AVM cell body as a landmark. In these conditions, the variance of AVM is also higher after starvation in the L3 stage compared to the control (*F*=22.6, *p*<10^−5^) (Figure S1A). Out of 200 animals in each condition, we only observed three animals with defects in axon (defasciculation, hook or extra branching) in the control population and also three after starvation. Axon formation is thus robust to early starvation and differences in cell position. Nonetheless, we found that AVM cell body (Figure S1C) or projection (Figure S1D) was more frequently in the vicinity of PLM projection after starvation (*LR X*^2^=16.8, *p*<10^−4^). An overlap between PLM projection and ALM has been previously observed in *sax-1* and *sax-2* mutants (Gallegos and Bargmann, 2004) but the overlapping between PLM and AVM was not discussed. More interestingly, we found that this close proximity (overlapping) is more prone to appear when AVM is posterior (*LR X*^2^=22.9, *p*<10^−5^) (Figure S1A).

### QR.pax final position is sensitive to body size

The end of QR.pa migration relies on the timing of expression of *mig-1* and not the position of the cell in the animal (Mentink et al., 2014). From this observation, we can predict that body size affects QR.pax final position: in a longer body, the cell should stop at the same time, i.e. at a more posterior position; conversely in a shorter body the cell should stop at a more anterior position (Figure 2A, left). Alternatively, body size variation may be fully compensated and the relative position of QR.pax remains unchanged at various body sizes (Figure 2A, right).

We thus tested whether body size affects QR.pax position. We selected a set of mutants in the N2 reference background with a smaller or a longer egg size and without general cell migration defect (Figure S2A). These include mutants in four genes as well as tetraploid animals with an overall larger body size (*C. elegans* is normally diploid (Nigon, 1951a)). *larp-1* encodes a La-related protein. A loss of function of this gene upregulates Ras-MAPK during oogenesis leading to smaller oocytes (Nykamp et al., 2008). The loss of function of *sma-1,* encoding a βH-spectrin, impairs embryonic elongation and leads to smaller body size (McKeown et al., 1998). *lon-1* encodes a cysteine-rich secretory protein (CRISP). Loss of function of this gene results in the upregulation of the TGF-β pathway, increasing body size by hypodermal endoreduplication, observed in adult stage (Morita et al., 2002). It also may affect egg length by physical compression in the gonad arm (Yamamoto and Kimura, 2017) but its effect on early larval development was not studied. *fln-1* encodes a filamin required for proper ovulation (Kovacevic and Cram, 2010). Nonetheless, how the loss of function allele *fln-1* (ok2611) affects egg and L1 length is unknown.

In these mutants, we measured QR.pax position and the pharynx-to-rectum (P- to-R) distance. The empirical measurements revealed that QR.pax final position and P-to-R distance are correlated, with longer animals having a more posterior QR.pax relative position (Figure 2B). This indicates that, as expected from the temporal regulation of its migration endpoint, QR.pax final positioning is sensitive to body size.

We established a mathematical model of the expected final position of QR.pax as a function of body size. The larva grows during the cell migration, as measured in Suppl. Table, which is taken into account in the model (see Methods). The model predicts the relationship between QR.pax position and body size at a given timepoint, in the absence of compensation (Figure 2C, red curve). Compared to the best-fit model without body size compensation, the empirical measurements show an effect on body size on QR.pax position that is smaller than that predicted by this simple model version, indicating the presence of a partial compensation mechanism.

We investigated two different scenarios of partial compensation to decipher the relationship between body size and QR.pax final position: an adaptation of the duration of migration of QR.p (but not QR.pa) to body size or an adaptation of the cell velocity to body size. In the first case, given that the temporal regulation of *mig-1* expression occurs at the end of the migration, we hypothesize that body size would affect the last part of the migration only, i.e. the short-range migration of QR.pa. Thus, the relative position of QR.p division should be similar between long and short animals. Measurements of the QR.p site of division (Rectum to QR.pp) and QR.pa site of division (QR.pp to QR.pa) did not support this hypothesis (Figure 2D), as the relative position of QR.p division is more anterior in small animals (*t* = −2.4, *p* = 0.02). In the second scenario, one can predict a mechanism through which the cell velocity is adapted to the body size, where the cell migrates faster in long animals. To estimate cell speed at different body sizes, we used the *sma-1* and *lon-1* mutants (the latter is close in body size to the wild-type N2; Figure 2B). We measured cell position at two timepoints during migration (t = 3h after hatching and t = 6h after hatching) and inferred cell velocity through the mathematical model, taking into account larval growth (Eq. 9). We found that the migration speed of the QR lineage from 3h to 6h after hatching was higher in *lon-1(e185); ayIs9* mutants (13.9 μm/hr) compared to *ayIs9 sma-1(e30)* mutants (9.3 μm/hr) (Figure 2E). These results are consistent with a partial compensation mechanism of body size acting on cell velocity. Indeed, incorporating these data into the model gives good agreement with the measurements, using no free parameters (Figure 2C, purple curve).

### Natural variation of QR.pax final position in wild isolates

QR.pax position has been studied only in the laboratory-adapted genetic background N2. The occurrence and span of natural variation for this trait was unknown. Above we detected natural variation among the four tested strains (Figure 1). We further measured QR.pax final position in a panel of 40 *C. elegans* strains (N2 and 39 wild isolates) that is representative of the genetic diversity of the species (Cook et al., 2017). We observed natural variation in QR.pax final position among the wild isolates (Figure 3A, *F* =18.9, *p*<10^−15^). The mean value of QR.pax final position in the N2 background is close to the grand mean in the species. Some isolates revealed a more anterior position (such as ECA36 and CB4932) and others a more posterior position (such as XZ1516 and JU830).

Considering the sensitivity of QR.pax final position to body size, we wondered whether natural variation in QR.pax position was explained at least partially by variation in body size. We first measured egg length as a proxy for L1 larval length that is easier to measure. However, when we measured larval lengths (P-to-R distances) at hatching and 6 hours after hatching in a subset of 14 strains, we realized that they were surprisingly poorly correlated with egg length in natural isolates, suggesting natural variation in morphogenesis within the egg shell (Figure S3A,B). Note that the N2 laboratory strain has the longest P-to-R distance of the 14 tested isolates. Interestingly, we did not find any correlation between QR.pax final position and body size in this set of isolates, suggesting that body size did not explain the variation in QR.pax final position and was thus compensated during evolution of the *C. elegans* species (Figures 3C and S3D, E). We did not find any correlation between QR.pax final position and the absolute value of the sampling latitude (*t* = −0.94, *p*=0.35).

## DISCUSSION

### Variance in final cell position

To inquire whether the temporally-regulated position of QR.pa daughters is sensitive to various perturbations, we used different types of metrics. I) For most measurements, we used a scale that takes as landmarks the lateral epidermal seam cells, as widely used in studies of QR/QL positioning (Coudreuse et al., 2006; Harris et al., 1996; Mentink et al., 2014; Whangbo and Kenyon, 1999). Although nematodes are not segmented, their organization is formed of repeats along the antero-posterior axis, particularly strikingly of these lateral epidermal seam cells. In our scale, each such repeat along the antero-posterior is further divided in four units, with a total of 27 units between the base of the pharynx to the rectum (Figure 1B). Note that QR is born prior to hatching as the sister of the lateral epidermal seam cell V5R and therefore its starting point is always the same on this scale (Sulston et al., 1983; Hedgecock et al., 1987). II) In some cases, we measured cell position in micrometers from the base of the pharynx or from the rectum (e.g. Figure 2D,E). III) When we varied body size, we scaled the micrometer measurements to a body length measure (e.g. Figure 2D).

We find that the variance in QR.pax position, as well as the observed range, stay within the length of a repeat along body length (4 of our scale units). The range encompasses about half a repeat length (2 scale units; Figure 2). However, after starvation in the L1 stage, the observed range encompasses over a repeat length. Using better known animals as a comparison, this range of QR.pax sensitivity to stochastic noise in the starvation treatment would correspond to the antero-posterior displacement of an insect appendage to the next segment, which would be considered a dramatic homeotic transformation. Note that in this starvation experiment, we only scored animals where development of seam cells and the QR lineage was not strongly retarded (see Methods) and thus the range may be even greater if all animals were considered. The increase in variance may be explained by the fact that after a long L1 starvation period, the growth speed of the worms is affected and development is delayed (Lee et al., 2012). This delay can potentially affect the timing of V seam cell division (Olmedo et al., 2020) as well as the intrinsic dynamics of QR migration. Importantly, the fact that BDU, ALM, CAN and HSN positions are not affected by early starvation also suggests that the relative scale used to measure the final position is not impaired by this treatment.

To some extent, displacements in the final position of the cell body can be compensated. For example in neurons, axon growth can be modulated according to the distance to the target by cell extrinsic determinants (Hekimi and Kershaw, 1993). Nonetheless in other cases, a shift in the position of neuronal cell bodies can drastically alter their fate and morphology (Martineau et al., 2018).

We wondered whether changes in the final position of QR.pax lead to differences in downstream neuronal phenotypes. We observed that posteriorly shifted AVM tends to be in the vicinity of the PLM projection after early starvation. We do not know whether a synaptic connection is established between these two mechanosensory neurons and whether this may be (counter)selected.

### A model without free parameters suggests partial compensation of body size

We observed that cells migrate farther in the animal relative to landmarks when body size is small, and less far when it is large. This observation is qualitatively consistent with the fact that these cells stop after a certain amount of time: if the speed is constant, they will migrate a constant distance, which will be larger relative to a smaller body. However, quantitatively, the degree of this effect was observed to be less than predicted by the model, even for the best-fit value of the constant speed. Indeed, in absolute values (microns), cells migrate farther when body size is large, and less far when body size is short (Figure S3B). We therefore hypothesized that partial compensation may be acting through a change in cell speed as a function of body size, i.e. if QR and its progeny migrate faster at larger body size. To test this hypothesis, we measured cell speed in *sma-1* versus *lon-1* mutants (Figure 2D) and using the model, we indeed found that the partial compensation of body size did operate through a change in cell speed. The improved model provided a quantitatively accurate description of the measurements. Moreover, given the known inputs (cell start position, cell start and stop time, cell velocity in two mutants, and larval growth speed), the improved model succeeded with no fit parameters.

Our mathematical model is minimal in its construction yet quantitatively accounts for the observations. Some simplifying assumptions are supported by the data, for example the observation that all larvae, independent of mutant strain, grow a constant amount in 6 hours. We have checked other assumptions explicitly; for example, we find that the results are negligibly changed if the cell accelerates and decelerates instead of instantaneously starting and stopping its migration. This result also suggests that our results would be robust to details such as temporal variations in cell speed or pauses due to cell division.

Possible mechanisms of cell speed dependence on body size could be the following. First, migration speed could be increased at larger cell size. Nonetheless, evidence from the literature tends to associate a negative correlation between cell size and cell velocity in vitro under adhesive conditions (Leal-Egaña et al., 2017; Hennig et al., 2020). Second, body size could affect extracellular matrix density, such that larger cells secrete a less tight matrix, resulting in faster net migration speed, as in the *emb-9* matrix collagen mutant (Kawano et al., 2009; Mentink et al., 2014). Third, body size could affect the properties of the Wnt gradient influencing QR migration; this could operate if for instance a larger body size resulted in stronger Wnt concentrations at a given relative position, resulting in faster cell speed (Mentink et al. 2014). Finally, it is possible that the cell speed modulation mechanism depends upon the specific mutants we used for these assays.

### Natural variation

All previous studies of QR migration were performed in the laboratory-modified N2 background of *C. elegans*. Here we uncover variation in the final position of QR.paa and QR.pap when exploring a representative set of *C. elegans* wild isolates. The variance in QR.pax position appears similar in all isolates but the mean position differs from about 1/4 of a body repeat (1 on our measurement scale) from CB4932 to XZ1516 (Figure 3A) in standard laboratory conditions. When considering different individuals, QR.pax position may vary in a range of almost a full body repeat among different individuals of different wild genotypes (3.75-7.5 on our scale; see Figure 3B for such representative animals).

Body size variation in the L1 stage is substantial among wild isolates, and almost covers the range of the body size mutants we used, excluding the tetraploid animals. However, we did not find a correlation between QR.pax body position and L1 body size in the subset of wild isolates we used for this analysis. This may be due to a lack of power or may suggest the presence of an evolutionary compensation in QR.pax position relative to body size variation. QR velocity and its migration dynamics can be affected in a different manner in each isolate. In any case, we conclude that natural variation in QR.pax position is not fully explained by body size variation. The available natural variation in QR.pax position will allow us in the future to analyze the genetic basis for the observed natural variation in the final position of a long-range migrating cell.

## Acknowledgements

We thank Hendrik Korswagen and his laboratory for discussions, as well as Michel Labouesse. We thank Hendrik Korswagen, Erik Schild and Joao Picao Osorio for comments on the manuscript. Some N2 mutant strains were provided by the CGC, which is funded by NIH Office of Research Infrastructure Programs (P40 OD010440). We thank Wormbase. This work was funded by a Collaborative Grant from the Human Frontier Science Program (RGP0030/2016).

The authors declare that they have no conflict of interests.

---

**Supplementary figure 1: A)** PLM projection overlaps with posteriorly shifted AVM after early starvation. Color dots represent the relative position of AVM for each animal, where its cell body or projection overlaps with PLM projection (light blue) or not (pink). Black dots and error bars represent the mean and SD of the two conditions (starved and non-starved); n=100 per replicate and condition, two replicates merged. The color dots associated with colored error bars represent the mean and SD of the overlapping (red) and non-overlapping (blue) distribution. **B-D)** Fluorescent pictures of animals at the L3 stage (GOU174 strain with GFP reporter in mechanosensory neurons). Without early starvation, PLM projection stops posteriorly to ALM and AVM cell body **(B)**. After early developmental arrest, the projection of PLM can overlap with AVM cell body **(C)** or AVM projection **(D)**.

**Supplementary figure 2: A)** Egg size measurements of mutants chosen to potentially affect egg size. WT: wild type, i.e. N2 reference background for all mutant lines. Black dots and error bars represent the mean and SD per genotype(n>20 per strain). Two-sided t-test comparison against WT with Bonferroni correction of the *p*-values. **B)** QR.pax final position in mutants showing a different egg size length. Grey dots represent QR.pax final position of each animal; black dots and error bars represent the mean and SD per genotype, respectively (n>40 per genotype). Two-sided t-test comparison against WT with Bonferroni correction of the *p-*values. ***: *p*<0.001, *: *p*<0.05. **C-D)** Relationships between egg length, pharynx-to-rectum (P-to-R) distance at 0 and 6h after hatching and QR.pax final position in N2 (green) and its size mutants (pink) Black dots and error bars represent the mean and confidence intervals (95%) for each genotype. The black line represents the regression line from the data, with the associated equation. Each color dot represents the mean position after random subsampling of 20 animals and the grey area represents regression lines after each iteration of subsampling (1000 iterations). *P*’ and *R^2^_adj_’* are the median of *P* and *R^2^_adj_’* after 1000 iterations.

**Supplementary figure 3:** Relationships between egg length, pharynx-to-rectum (P-to-R) distance at 0 and 6h after hatching and QR.pax final position in N2 (green) and a subset of wild isolates (blue). Black dots and error bars represent the mean and confidence intervals (95%) for each genotype. The black line represents the regression line from the data, with the associated equation. Each color dot represents the mean position after random subsampling of 20 animals and the grey area represents regression lines after each iteration of subsampling (1000 iterations). *P*’ and *R^2^_adj_’* are the median of *P* and *R^2^_adj_’* after 1000 iterations.

## Supplementary Table

Raw data and summary of: QR.pax, BDU, ALM, CAN and HSN final position measurements for the robustness to noise (sheet 1), QR.pax final position measurement for the robustness to temperature (2a) and starvation (2b), AVM distance, overlap and defect count (3),natural variation in the panel of 40 strains (4), egg length measurements (5), Pharynx to rectum distance at hatching and 6h after hatching (6), QR.pax final position in N2 mutants (7), QR.p and QR.pa site of divisions (8) and cell velocity (9).

